# Proteomic Profiles of Growth-Related Proteins in Red Sokoto and West African Dwarf Goats

**DOI:** 10.64898/2026.01.23.701243

**Authors:** O. Babatunde, O.A. Akintunde, B.A. Ajayi

## Abstract

Proteomic profiling provides a framework for describing breed-specific growth, physiological and adaptive mechanisms in livestock. This study compared the plasma proteomic profiles of Red Sokoto (RS) and West African Dwarf (WAD) goats, with emphasis on growth-related proteins. Plasma samples from 20 goats (RS = 10; WAD = 10) collected across three locations in southwestern Nigeria were analysed using liquid chromatography–mass spectrometry (LC–MS). A total of 66 plasma proteins were identified in RS goats and 59 in WAD goats, of which 14 were associated with growth regulation. Distinct breed-specific expression patterns were evident. RS goats exhibited higher abundance of fibronectin and calmodulin, indicating enhanced tissue remodelling, muscle development, and calcium-mediated signaling. In contrast, WAD goats showed relatively higher expression of key metabolic and endocrine regulators, including insulin, leptin, ghrelin, glucagon, adiponectin, epidermal growth factor, erythropoietin, and thrombopoietin, reflecting greater metabolic efficiency and adaptive resilience. Gene ontology enrichment analysis revealed marked functional divergence between breeds: RS goats demonstrated stronger enrichment of GO terms related to signal transduction efficiency, cell–matrix adhesion, calcium ion binding, and growth-related morphogenetic processes, whereas WAD goats showed enrichment of GO categories associated with energy metabolism, stress adaptation, catabolic regulation, and hematopoietic support. These findings indicate that breed differences in growth potential are driven more by pathway efficiency and functional integration than hormone abundance. Plasma proteomic and GO-based functional profiles highlight coordinated anabolic and structural growth regulation in RS goats and a resilience-oriented metabolic strategy in WAD goats, with important implications for breed-specific selection, conservation, and sustainable goat production systems.

## Introduction

Indigenous goat breeds play critical roles in food security, income generation, and livelihood sustainability in Nigeria, particularly within low-input production systems. Among indigenous Nigerian goats, the Red Sokoto (RS) and West African Dwarf (WAD) goats exhibit distinct phenotypic divergence in body size, growth rate and environmental adaptability (Yakubu et al., 2011). Red Sokoto goats are widely valued for superior body conformation, meat yield, and high-quality leather (Yakubu & Mohammed, 2012), whereas WAD goats are distinguished by exceptional resilience, feed efficiency, and tolerance to nutritional and climatic stressors common in humid tropical environments (Daramola & Adeloye, 2009).

Understanding the molecular mechanisms underlying these breed-specific traits is essential for improving productivity, conservation and breeding strategies. Proteomics provides an approach for investigating functional biological processes by directly characterizing expressed proteins that mediate metabolism, growth, immune responses, and environmental adaptation (Al-Amrani et al., 2021; Bourganou et al., 2025). Plasma proteomics is particularly informative, as circulating proteins integrate signals from multiple tissues and reflect the physiological status of the organism under prevailing environmental and genetic influences (Geyer et al., 2024).

Despite the economic and adaptive importance of indigenous goat breeds, information on breed-specific plasma proteomic profile, particularly for growth-related proteins (GRP), in Nigerian goats is limited. Most previous studies have relied on phenotypic, morphometric or genetic markers, offering limited insight into the functional biochemical pathways driving breed differences (Akintunde et al., 2024; Babatunde et al., 2025). Therefore, this study aimed to characterize and compare the plasma proteomic profiles of Red Sokoto and West African Dwarf goats using liquid chromatography–mass spectrometry (LC–MS), with specific emphasis on growth- and tissue development–related proteins.

## Materials and Methods

The study was conducted at selected goat-rearing locations in Osun, Ondo, and Oyo States in southwestern Nigeria. Prior to sample collection, visits were made to identify suitable animals and to obtain informed consent from goat owners. All animal handling and blood sampling procedures were carried out by trained personnel under the supervision of certified animal scientists and were performed in accordance with institutional and national guidelines for the care and use of animals in research (UNIOSUNHREC 2025/014B).

A total of 20 apparently healthy adult goats comprising Red Sokoto (RS; n = 10) and West African Dwarf (WAD; n = 10) breeds were randomly selected for the study. Sampling was conducted across three locations as shown in Table 1: Osogbo, Osun State (4 RS and 4 WAD); Owena, Ifedore Local Government Area, Ondo State (3 RS and 3 WAD); and Ajaawa, Ogo-Oluwa Local Government Area, Oyo State (3 RS and 3 WAD). Only newly arrived RS goats from northern Nigeria, where they were raised, and that had been in the area for no more than two days were used for the study. Approximately 2 mL of blood was aseptically collected from the jugular vein of each goat using sterile disposable syringes and transferred into pre-labelled EDTA microtainer tubes. The samples were gently inverted several times to ensure thorough mixing with the anticoagulant and were immediately placed in ice-packed containers. Blood samples were centrifuged at 3,000 × g for 10 min at 4 °C to separate plasma. The plasma fraction was carefully aliquoted into sterile microcentrifuge tubes. The samples were assembled immediately after collection and taken to Ibadan without delay, where they were transferred into a cryogenic flask and conveyed at approximately −80 °C to the Molecular Laboratory, Bayero University, Kano. Samples were received within 48 hours of collection and stored at −20 °C until proteomic analysis.

**Table 1:** Sampling locations and distribution of Red Sokoto (RS) and West African Dwarf (WAD) goats used for plasma proteomic analysis.

| State | Location | RS (n) | WAD (n) | Total (n) |
| --- | --- | --- | --- | --- |
| Osun | Osogbo | 4 | 4 | 8 |
| Ondo | Owena (Ifedore LGA) | 3 | 3 | 6 |
| Oyo | Ajaawa (Ogo-Oluwa LGA) | 3 | 3 | 6 |
| **Total** |  | **10** | **10** | **20** |

*RS = Red Sokoto goats; WAD = West African Dwarf goats; LGA = Local Government Area.*

For proteomic analysis, plasma samples were diluted (1:4, v/v) in 8 M urea prepared in 50 mM Tris-HCl buffer (pH 8.0) to promote protein denaturation prior to enzymatic digestion. Proteins were enzymatically digested into peptides using trypsin under controlled conditions. Following digestion, solid-phase extraction (SPE) was employed to remove impurities and concentrate peptides prior to mass spectrometric analysis. Peptide samples (50 µL) were injected into a Shimadzu liquid chromatography–mass spectrometry (LC–MS) system equipped with a triple quadrupole mass analyzer. Mass spectrometry parameters, including ionization mode, collision energy, and scan range, were optimized for plasma protein detection. Acquired raw data were processed using ProteoWizard software for protein identification and quantification (Kessner et al., 2008; Chambers et al., 2012). Identified proteins were annotated and functionally classified using UniProt and the National Center for Biotechnology Information (NCBI) databases (UniProt Consortium, 2023).

Protein concentrations (µg/µL) and corresponding retention times were obtained for all identified proteins. The GRP were extracted based on functional annotation and visualized using heatmaps and bar charts generated with GraphPad Prism version 10. Gene ontology information was used to support functional interpretation of the identified proteins.

## Results

Proteomic profiling of plasma samples identified 66 proteins in RS goats and 59 proteins in WAD goats. Functional annotation and Gene Ontology (GO) classification identified 14 proteins as growth-related (Table 2). Heatmap analysis revealed clear breed-specific protein abundance patterns (Fig. 1). Structural and signaling proteins, notably fibronectin and calmodulin, were detected at high and moderate levels respectively in RS goats but were below the detection threshold in WAD goats. Most endocrine growth regulators such as insulin, leptin, ghrelin, glucagon, adiponectin, vascular endothelial growth factor (VEGF), and epidermal growth factor (EGF) were present at generally low abundance in both breeds. However, erythropoietin (EPO) and thrombopoietin exhibited slightly higher abundance in WAD goats. Comparative bar-chart analysis with GraphPad prism confirmed breed-associated differences in GRP (Fig. 2a & 2b) WAD goats consistently showed relatively higher plasma levels of insulin, leptin, ghrelin, glucagon, adiponectin, EGF, EPO, and thrombopoietin than RS goats, whereas VEGF abundance was comparable between breeds. In contrast, proteins associated with extracellular matrix organization and intracellular signaling displayed opposite trends. Fibronectin and calmodulin were substantially more abundant in RS goats, indicating enhanced structural and calcium-mediated signaling capacity. Matrix metalloproteinase-9 (MMP-9) was detected at low levels in both breeds, with slightly higher abundance observed in WAD goats.

**Table 2:**
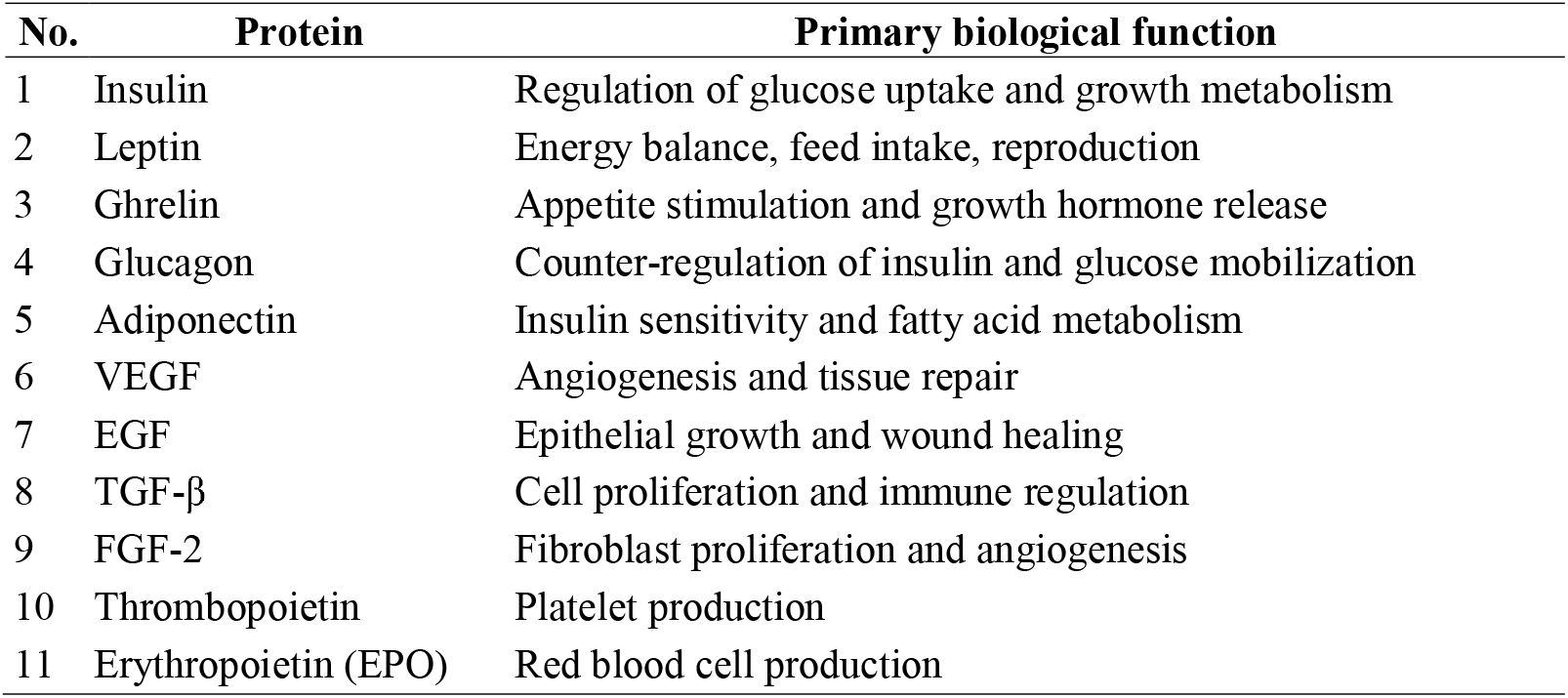
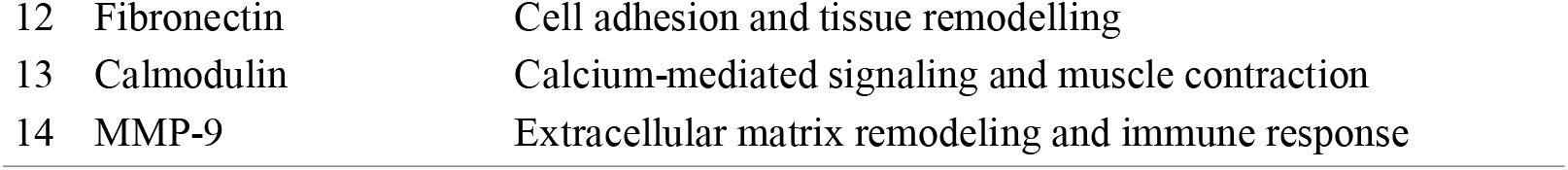
Growth-related plasma proteins identified in Red Sokoto and West African Dwarf goats and their biological functions.

| No. | Protein | Primary biological function |
| --- | --- | --- |
| 1 | Insulin | Regulation of glucose uptake and growth metabolism |
| 2 | Leptin | Energy balance, feed intake, reproduction |
| 3 | Ghrelin | Appetite stimulation and growth hormone release |
| 4 | Glucagon | Counter-regulation of insulin and glucose mobilization |
| 5 | Adiponectin | Insulin sensitivity and fatty acid metabolism |
| 6 | VEGF | Angiogenesis and tissue repair |
| 7 | EGF | Epithelial growth and wound healing |
| 8 | TGF-β | Cell proliferation and immune regulation |
| 9 | FGF-2 | Fibroblast proliferation and angiogenesis |
| 10 | Thrombopoietin | Platelet production |
| 11 | Erythropoietin (EPO) | Red blood cell production |
| 12 | Fibronectin | Cell adhesion and tissue remodelling |
| 13 | Calmodulin | Calcium-mediated signaling and muscle contraction |
| 14 | MMP-9 | Extracellular matrix remodeling and immune response |

**Table 3:** Relative abundance patterns of growth-related plasma proteins in Red Sokoto and West African Dwarf goats based on heatmap analysis.

| Protein | RS | WAD | Interpretation |
| --- | --- | --- | --- |
| Insulin | Very low | Very low | Uniformly low |
| Leptin | Very low | Very low | Uniformly low |
| Ghrelin | Very low | Very low | Uniformly low |
| Glucagon | Very low | Very low | Uniformly low |
| Adiponectin | Very low | Very low | Uniformly low |
| VEGF | Very low | Very low | Uniformly low |
| EGF | Very low | Very low | Uniformly low |
| TGF-β | Missing | Missing | Not analysed |
| FGF-2 | Missing | Missing | Not analysed |
| Thrombopoietin | Very low | Low | Slightly higher in WAD |
| EPO | Low | Low | Uniformly low |
| Fibronectin | High | Missing | Highly expressed in RS |
| Calmodulin | Moderate–high | Missing | More abundant in RS |
| MMP-9 | Low | Low | Uniformly low |

*Missing = protein whose level of detection is insignificant.*

**Figure 1:**
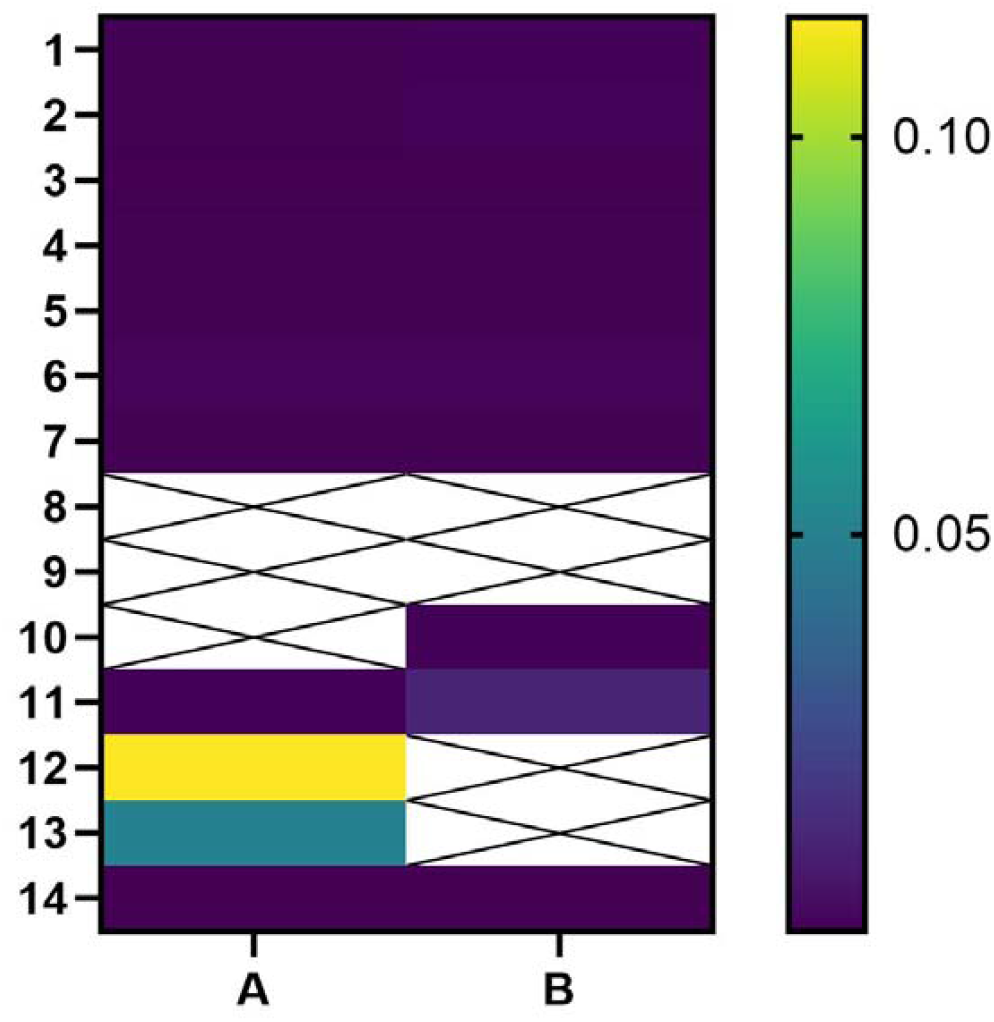
Heatmap showing relative abundance of growth-related plasma proteins in Red Sokoto (A) and West African Dwarf (B) goats. Yellow-green indicates high relative abundance, dark purple indicates low or undetectable abundance, and white (X) represents missing or excluded data. RS goats show higher abundance of fibronectin and calmodulin, whereas most metabolic hormones exhibit uniformly low abundance in both breeds.

**Figure 2a.**
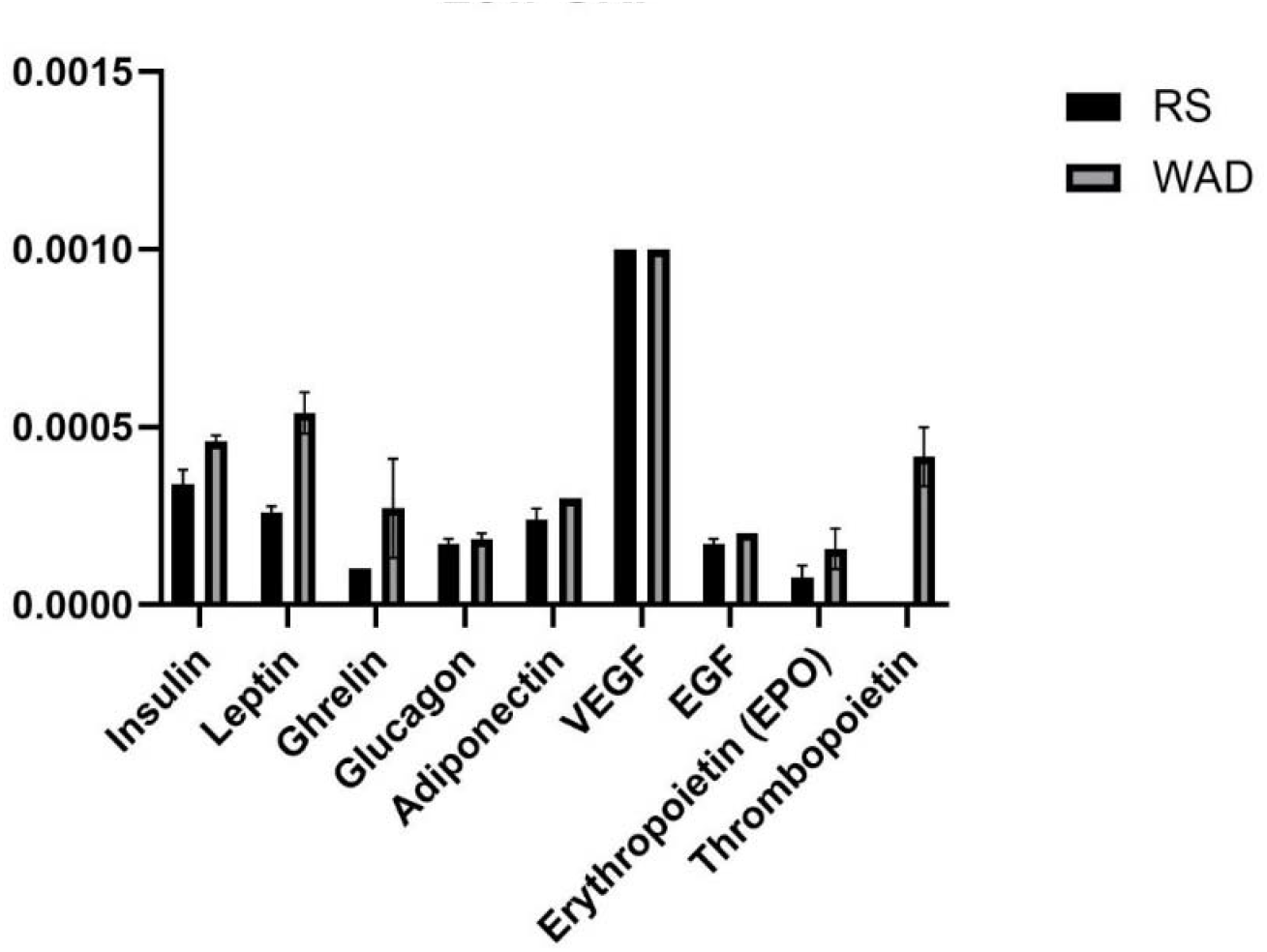
Comparative concentrations of growth-related proteins in Red Sokoto and West African Dwarf goats expressed in µg/µL. Bar charts illustrate quantified plasma concentrations (µg/µL) of insulin, leptin, ghrelin, glucagon, adiponectin, VEGF, EGF, erythropoietin, and thrombopoietin. WAD goats generally exhibit higher levels of metabolic and hormonal regulators, indicating enhanced metabolic adaptability.

**Figure 2b.**
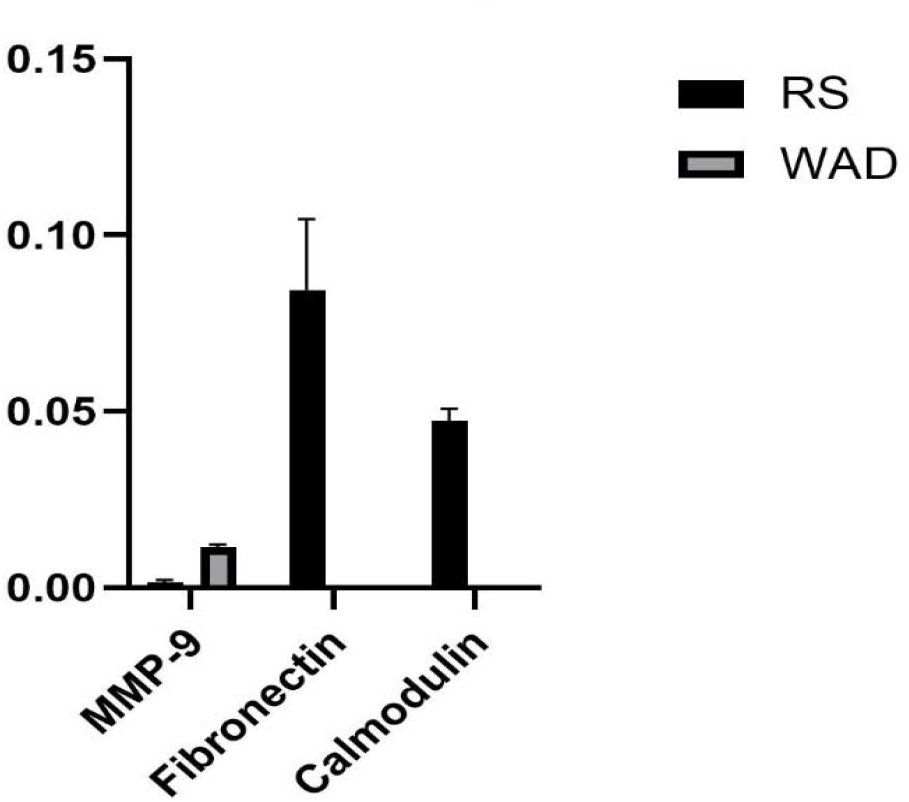
Comparative concentrations of growth-related proteins in Red Sokoto and West African Dwarf goats expressed in µg/µL

GO enrichment analysis of growth-related proteins revealed marked functional divergence between breeds. Although WAD goats exhibited higher circulating levels of insulin, leptin, ghrelin, glucagon, and adipokines, associated GO terms were predominantly linked to metabolic regulation, stress adaptation, and energy conservation. In contrast, RS goats showed stronger enrichment of GO terms related to signal transduction efficiency, calcium-mediated signaling, cell–matrix adhesion, and tissue morphogenesis, despite lower hormone abundance. GO terms associated with insulin-, leptin-, and ghrelin-related pathways indicated greater enrichment of PI3K–Akt and MAPK signaling, cellular proliferation, and GH/IGF-axis responsiveness in RS goats, whereas WAD goats showed signatures consistent with reduced downstream signaling efficiency. Structural and signaling proteins further differentiated the breeds, with RS goats showing enrichment for extracellular matrix organization and calcium ion binding, and WAD goats showing higher representation of catabolic, remodeling, and hematopoietic processes, including glucagon, adiponectin, MMP-9, EPO, and thrombopoietin. VEGF-associated GO terms were similar between breeds.

### Expression of structural and signalling proteins in Red Sokoto and West African Dwarf goats

Relative abundance of MMP-9, fibronectin, and calmodulin demonstrates enhanced tissue remodelling and calcium-mediated signalling in RS goats compared with WAD goats.

## Discussion

The plasma proteomic profiles of West African Dwarf (WAD) and Red Sokoto (RS) goats reveal distinct physiological strategies underlying breed-specific growth performance. In WAD goats, the relative enrichment of metabolic and endocrine regulators suggests a system optimized for metabolic flexibility and efficient energy utilization. Hormones such as insulin and glucagon coordinate glucose homeostasis, while leptin and ghrelin regulate appetite, energy balance and nutrient partitioning (Chilliard et al., 2005; Wren et al., 2001; Dimitriadis et al., 2021; Venugopal et al., 2023). The higher abundance of these regulators in WAD goats is consistent with their ability to survive and remain productive under nutrient-limited and environmentally challenging tropical conditions. Adiponectin further supports this profile by enhancing insulin sensitivity and promoting fatty acid oxidation, thereby favoring energy efficiency and metabolic resilience (Choi et al., 2020). Elevated erythropoietin (EPO) may additionally reflect adaptive support for oxygen transport and aerobic metabolism under environmental or thermal stress, although direct physiological measurements would be required to confirm this association (Schoener & Borger, 2024).

In contrast, RS goats exhibited a plasma proteomic signature dominated by proteins associated with tissue organization and intracellular signaling. The higher abundance of fibronectin, a key extracellular matrix glycoprotein involved in cell adhesion, tissue remodeling, and muscle development, is indicative of enhanced anabolic and structural activity (Stoffels et al., 2013; Grzelkowska-Kowalczyk, 2016). Similarly, elevated calmodulin abundance suggests active calcium-mediated signaling pathways essential for muscle contraction, cellular regulation, and growth (Tansey et al., 1994; Kaleka et al., 2012). These molecular features align with the recognized growth potential, body conformation, and carcass characteristics of RS goats and support their suitability for meat-oriented production systems. Matrix metalloproteinase-9 (MMP-9), which participates in extracellular matrix turnover and immune-related tissue remodeling, was detected at low levels in both breeds but showed slightly higher abundance in WAD goats. This pattern may reflect a greater capacity for tissue adaptability and stress responsiveness, consistent with the ecological versatility of WAD goats (Li et al., 2016; He et al., 2023). However, as the present analysis is based on plasma proteomics, these findings likely represent systemic regulatory signals rather than localized tissue-specific activity.

Beyond individual proteins, the broader proteomic patterns observed between RS and WAD goats reflect fundamental differences in growth regulation. RS goats, which exhibit superior growth and larger body size, showed enrichment of proteins involved in amino acid, glucose, and lipid metabolism, whereas WAD goats displayed higher abundance of acute-phase and structure-related proteins linked to immune function and stress tolerance. Similar trends have been reported in cattle and other ruminants, where faster-growing and feed-efficient animals exhibit upregulation of metabolic and mitochondrial proteins, while slower-growing or less efficient animals show greater expression of extracellular matrix and stress-related proteins (Idowu et al., 2024). Evidence from transcriptomic and metabolomic studies further supports the central role of downstream metabolic efficiency in growth regulation. In goats, enhanced growth has been associated with upregulation of genes involved in lipid transport, fatty acid metabolism, and mitochondrial energy production, alongside activation of AMPK and Toll-like receptor signaling pathways (Wang et al., 2023). Metabolomic analyses similarly indicate that growth-related biomarkers are dominated by lipid classes and organic acids reflecting energy mobilization and anabolic capacity, rather than endocrine hormone concentrations (Li et al., 2024). Comparable findings in cattle and sheep demonstrate that metabolic and signaling proteins correlate more strongly with growth rate and feed efficiency than circulating growth hormone or insulin-like growth factor-1 (IGF-1), which often show limited discriminatory power between growth phenotypes (Idowu et al., 2024; Liu et al., 2025).

Although growth hormone and IGF-1 are essential for postnatal growth, accumulating evidence indicates that hormone concentration alone is insufficient to explain inter-breed variation. Positive associations between IGF-1 and body weight have been reported in several goat breeds, including Saanen and Angora kids, with IGF-1 levels influenced by environmental factors such as photoperiod and temperature (Pehlivan, 2019). However, functional studies show that downstream modulators of hormone action can exert stronger effects on growth than hormone abundance itself. Notably, IGF binding protein 2 (IGFBP2) negatively regulates muscle growth in goats by modulating TGF-β signaling and mitochondrial biogenesis, with genetic variation in IGFBP2 associated with reduced growth performance despite normal endocrine profiles (Liu et al., 2025). These findings align closely with the present proteomic results and reinforce the importance of signaling efficiency and metabolic responsiveness in determining growth outcomes.

The contrasting proteomic profiles of RS and WAD goats also reflect adaptive trade-offs between growth and resilience. WAD goats are widely recognized for exceptional tolerance to trypanosomiasis, gastrointestinal nematodes, and environmental stressors, traits that are genetically and immunologically mediated (Chiejina & Behnke, 2011). Sustained investment in immune surveillance and stress-response pathways likely diverts nutrients and energy away from tissue accretion, constraining growth. This trade-off is well documented across livestock systems, where animals selected for robustness and survivability typically exhibit lower growth potential due to higher maintenance and defense costs (Gaughan et al., 2019). Reviews across ruminant species further indicate that low-growth animals often maintain superior homeostasis under nutritional and thermal stress, whereas high-producing breeds are more vulnerable to physiological disruption (Gaughan et al., 2019; Silanikove, 2000). Heat stress and immune activation provide clear mechanistic examples of this balance. High-growth ruminants generate greater metabolic heat and show increased susceptibility to thermal stress, whereas smaller or indigenous breeds demonstrate superior thermoregulation at the expense of growth rate (Gaughan et al., 2019; Sejian et al., 2018). Immune challenges similarly suppress anabolic pathways through cytokine-mediated inhibition of appetite and IGF-1 signaling, constraining growth during disease pressure (Lochmiller & Deerenberg, 2000; Doeschl-Wilson et al., 2009). In addition, early sexual maturity and high reproductive frequency in indigenous goats impose further energetic demands that limit somatic growth (Silanikove, 2000). The higher abundance of immune- and structure-related proteins in WAD goats observed in this study is therefore consistent with a survival-oriented growth strategy.

These findings demonstrate that growth variation between Nigerian WAD and RS goat breeds is governed by coordinated molecular regulation involving metabolic enzymes, signaling mediators, and structural proteins. Plasma proteomics, supported by transcriptomic, metabolomic, and hormone-based evidence, highlights the primacy of downstream cellular processes in shaping growth performance. Red Sokoto goats appear optimized for efficient anabolic metabolism and tissue accretion under favorable conditions, whereas West African Dwarf goats prioritize resilience and survival in challenging environments, even at the cost of reduced body size. These breed-specific strategies mirror patterns observed across African and non-African ruminant breeds and underscore the value of integrative molecular approaches for understanding growth, adaptation, and productivity in indigenous livestock populations (Idowu et al., 2024; Gaughan et al., 2019).

## Conclusion

The proteomic and gene ontology evidence generated in this study provides a functional basis for breed-targeted goat improvement in Nigeria. The coordinated enrichment of structural and signalling pathways related to tissue development in Red Sokoto goats supports their prioritization for growth- and meat-oriented selection, particularly in systems where carcass evaluation is limited. Conversely, the dominance of metabolic and adaptive regulatory pathways in West African Dwarf goats confirms their value as a genetic resource for resilience, metabolic efficiency, and survival under low-input tropical conditions. The findings demonstrate that effective selection for growth should focus on pathway efficiency and functional protein markers rather than hormone abundance alone. This can complement conventional phenotypic selection, guide structured crossbreeding strategies, and support the development of sustainable, climate-resilient goat breeding programmes.

## Declarations

### Ethical Approval Certificate

The experimental procedures of this study were approved by the UNIOSUN HREC Committee of Osun State University, Osun, Nigeria (26^th^ September, 2025; UNIOSUNHREC 2025/014B).

## Author Contribution Statement

O. Babatunde: Data collection, investigation, formal analysis, and writing the original draft

A.O. Akintunde: Project administration, supervision, conceptualization, methodology, review and editing

B.A. Ajayi: Project administration, supervision, conceptualization, methodology, review and editing

## Acknowledgments

The following undergraduate students of the Department of Animal Science, Osun State University, Nigeria are acknowledged for their technical and material support in measuring the WAD and RS goats throughout the research work:

1. Aderogba Mubarak
2. Atoyebi Temidayo Oyewumi
3. Oyekanmi Omotayo Moses
4. Okeowo Lukman Adekunle
5. Ogunleye Toluwani Enoch
6. Akinlabi Kabirat Motunrayo
7. Fiyin Ebenezer Aremu

